# Projections between visual cortex and pulvinar nucleus in the rat

**DOI:** 10.1101/2020.01.27.921858

**Authors:** Leo R. Scholl, Andrzej T. Foik, David C. Lyon

## Abstract

The extrageniculate visual pathway, which carries visual information from the retina through the superficial layers of the superior colliculus and the pulvinar nucleus, is poorly understood. The pulvinar is thought to modulate information flow between cortical areas, and has been implicated in cognitive tasks like directing visually guided actions. In order to better understand the underlying circuitry, we performed retrograde injections of modified rabies virus in the visual cortex and pulvinar of the Long-Evans rat. We found a relatively small population of cells projecting to primary visual cortex (V1), compared to a much larger population projecting to higher visual cortex. Reciprocal corticothalamic projections showed a similar result, implying that pulvinar does not play as big a role in directly modulating V1 activity as previously thought.

---

Most visual information in primary visual cortex (V1) is delivered from the retina via the lateral geniculate nucleus (LGN; Jones, 1985). However, the extrageniculate visual pathway, which carries information from the retina through the superficial layers of the superior colliculus (SC) and the pulvinar nucleus (Kaas and Huerta, 1988; Stepniewska, 2003; Lyon et al., 2010), also makes major contributions to visual processing, having been implicated in gating visual cortex activity both in primates (Purushothaman et al., 2012; Zhou et al., 2016) and in rodents (Tohmi et al., 2014; Roth et al., 2015). In comparison to the LGN, less is known about the structure and function of the extrageniculate thalamic nuclei, yet these structures have a big impact on cognition and behavior. Early behavioral studies in primates identified cells in the pulvinar that are enhanced by shifts in gaze (Petersen et al., 1985; Robinson et al., 1986; Bender, 1982), leading many to think the pulvinar is involved in directing spatial attention. Modern theories of pulvinar’s role in attention include its driving of salience-based selection (Mizzi & Michael, 2014; Veale et al., 2016), guiding visual actions (Wilke et al., 2010; Zhou et al., 2017), and providing contextual information to visual cortex (Wilke et al., 2009; Roth et al., 2015; Jaramillo et al., 2019). Because of its reciprocal connections with so many visual areas and its behavioral role in both action and object vision, the pulvinar is a likely player in the modulation of information flow to the so-called dorsal and ventral visual streams (Kaas and Lyon, 2007), processing the guidance of action and object recognition, respectively (Goodale, 2005; 2013).

The rat pulvinar, also called the lateral posterior nucleus (LP), consists of three highly-conserved subregions based on cytoarchitecture and connectivity: caudomedial (LPcm), lateral (LPl), and rostromedial (LPrm) pulvinar (Takahashi, 1985; Nakamura et al., 2015; see Figure 1). Caudal LP receives input primarily from SC and pretectum (Takahashi, 1985; Mason & Groos, 1981; Shi & Davis, 2001) and sends projections mostly to temporal association cortex and postrhinal cortex (Nakamura et al., 2015; Shi & Davis, 2001), whereas rostral LP receives and sends most of its projections to visual cortex (Takahashi, 1985; Nakamura, 2015; Masterson, 2009; Bourassa & Deschenes, 1995). However, while some projections, such as retrosplenial cortex and amygdala projections from rostral LP have been studied in detail (Kamishina, et al., 2009), a quantitative analysis of rat pulvinar connectivity with visual cortex has not yet been made. The pulvinar nucleus is known to send projections broadly to visual cortex in several other species (Zhou et al., 2017), including mice (Tohmi et al., 2014; Bennett et al., 2019), gray squirrels (Robson and Hall, 1977), carnivores (Hutchins & Updyke, 1989; Mason, 1978) and primates (Benevento & Rezak, 1976; Asanuma et al., 1985; Adams et al., 2000). Nevertheless, it is difficult to interpret the role of the pulvinar on cortical activity without a more precise understanding of the relative weights of these connections, as well as the details of where in the pulvinar these projections originate.

**Figure 1.**
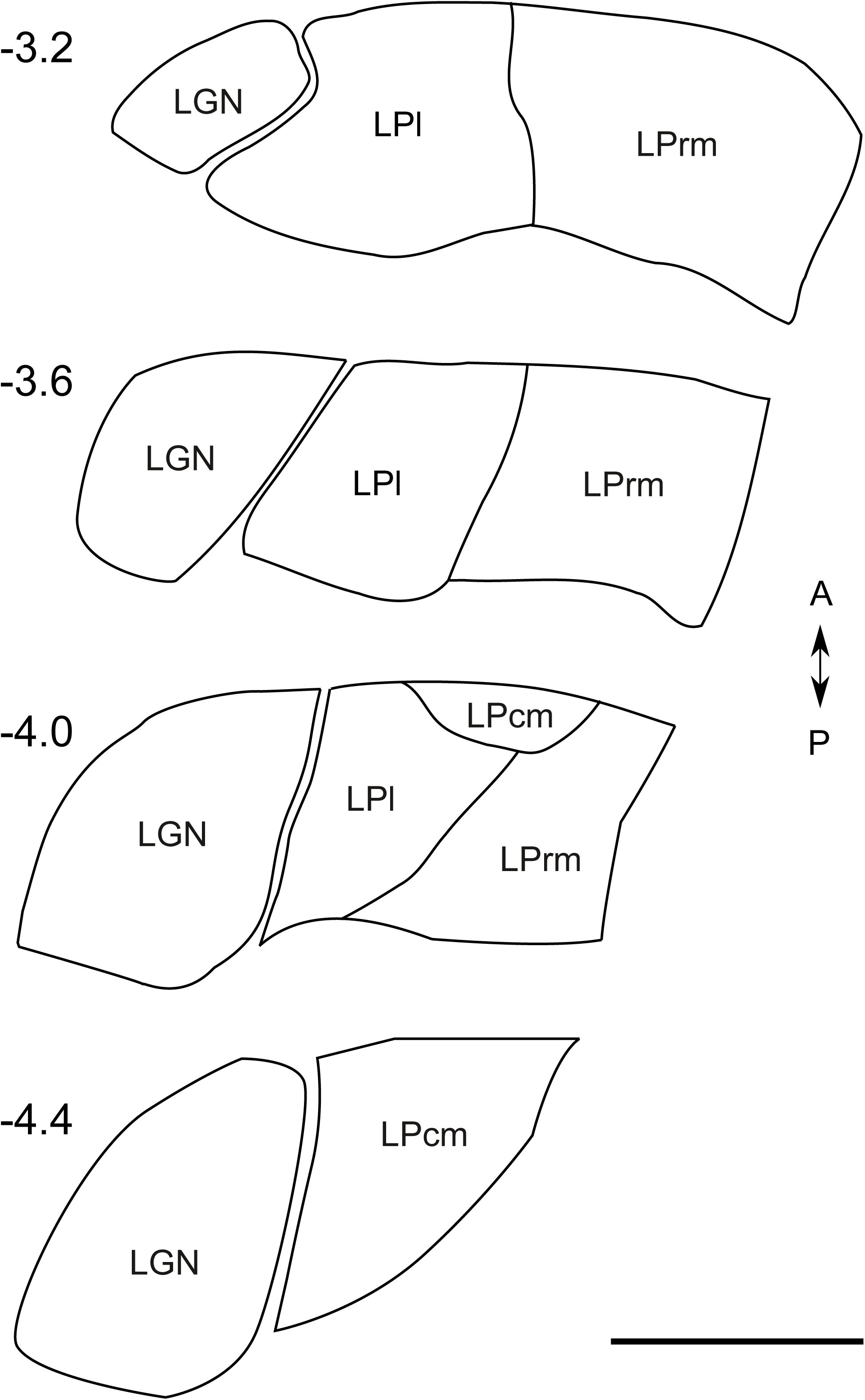
Schematic of rat lateral (LPl), rostromedial (LPrm) and caudomedial (LPcm) pulvinar subdivisions and the lateral geniculate nucleus (LGN) between −3.2 and −4.4 mm from bregma. Based on the cytoarchitectonic divisions by Nakamura (2015), conformed to the atlas by Paxinos and Watson (2013).

Of particular interest to us was whether pulvinar in rats projects more to V1 or V2. In rodents, V1 is homologous to primate V1, whereas higher visual cortex is called V2 and is subdivided into areas receiving retinotopically organized input from V1, including medial areas anteromedial (AM) and posteromedial (PM), as well as lateral areas anterolateral (AL) and lateromedial (LM) (Olavarria and Montero, 1984; Glickfeld, et al., 2014). Each of these areas likely receives some LP input, as has been demonstrated in mouse (Tohmi et al., 2014; Juavinett et al., 2019), but it remains unclear how contributions from pulvinar differ between higher visual cortex and V1. After lesions in mouse SC, higher visual cortex activity becomes more similar to activity in V1, suggesting that only higher visual cortex is affected by pulvinar (Tohmi et al., 2014). Yet mouse LP axon terminals do confer information to V1, as demonstrated by Roth et al. (2015), although it is unclear from how many LP cells these axons originate. To better understand the contributions of the pulvinar to visual cortical cells, a more complete map of its connectivity is needed.

To address these issues, we made injections of g-deleted rabies virus in V1 and higher visual cortex, as well as in lateral and rostromedial subdivisions of LP, to retrogradely label projection neurons. In this way we are able to determine whether or not there are quantitative differences in the thalamocortical and corticothalamic projections between V1 and higher visual cortex with the rat pulvinar.

## Methods

Injections of modified rabies virus were carried out in nine adult female Long-Evans rats in order to retrogradely label connected cells (Foik et al., 2018). All procedures were approved by the University of California, Irvine Institutional Animal Care and Use Committee and the Institutional Biosafety Committee, and followed the guidelines of the National Institutes of Health.

G-deleted rabies viruses (RV; Wickersham et al., 2007) modified with either green fluorescent protein (GFP) or mCherry transgenes (provided by the Callaway laboratory) were amplified and purified as described by Osakada & Callaway (2013). For each virus, BHK cells expressing rabies glycoprotein SADB19G (B7GG, provided by the Callaway laboratory) were infected with 1 µl of stock virus and maintained at 3% CO_2_ and 35°C for 5-6 d in order to produce viral supernatant (Figure 2a and 2b). The supernatants for each virus were subsequently used to infect five 150 mm dishes of the same cell line in order to amplify the viruses. Supernatants were collected twice during incubation at 3% CO_2_ and 35°C after 6 and 10 days, then passed through a 0.45 µm polyethersulfone filter, and transferred to an ultracentrifuge (rotor SW28, Beckman Coulter) for 2 h at 19,400 g and 4°C. Purified virus was resuspended in phosphate buffered saline (PBS) for 1 h at 4°C before 2% fetal bovine serum was added. Aliquots for injection were stored at −80°C. Titer was assessed by infecting HEK 293T cells (Sigma-Aldrich) with serial dilutions of modified virus, to ensure at least ∼1 x 10^9^ infectious units / mL was achieved. Figure 2 shows infected B7GG cells prior to amplification and infected 293T cells during titration.

**Figure 2.**
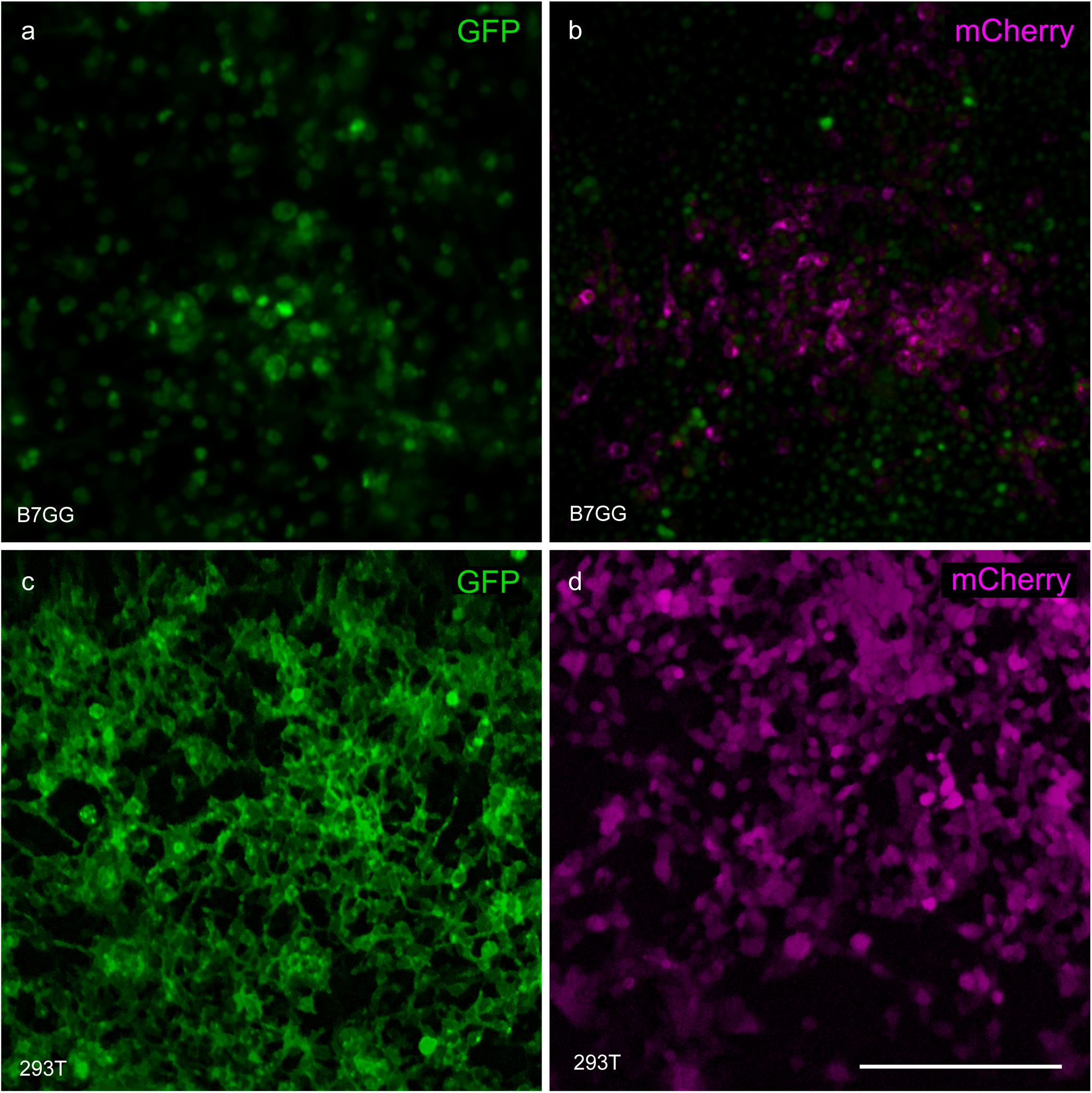
Amplification and titration of GFP and mCherry modified rabies viruses. B7GG cells in (a) and (b) express nuclear GFP in addition to the rabies viruses being amplified. 293T cells shown in (c) and (d) were used to verify expression of the viruses without the presence of cellular rabies glycoprotein and to quantify the virus titers. Scale bar equals 250 µm.

Prior to surgery, rats were initially anesthetized with 2% isoflurane in a mixture of 30% oxygen and 70% nitrous oxide, and maintained with 1 to 1.5% isoflurane in the same mixture. Using a stereotaxic apparatus, a craniotomy was performed to expose the caudal neocortex of one hemisphere. A glass micropipette was cut to approximately 20 µm in diameter, filled with rabies virus suspension, and lowered into the brain using a motorized microdrive to a depth of roughly 800 µm for cortical injections or 4,250 µm for LP injections. Stereotaxic coordinates were used to target each structure: for V1 injections, between −6 and −8 mm from bregma and between 3.75 and 4.25 mm from the midline; for medial V2 injections, −5.5 to −6.5 mm from bregma, 2.25 mm lateral; for lateral V2 injections, −6 to −7 mm from bregma, 5.5 mm lateral; for LPrm injections, −3.75 to −4 from bregma, 1.75 mm lateral; for LPl injections, −3.75 to −4 from bregma, 2.75 mm lateral. Viral suspensions were injected at a rate of approximately 1 µl/min using an adjustable regulator and pressures below 35 kPa, to a volume of no more than 1.2 µl per injection, as larger injection volumes can cause damage to surrounding tissue. To increase total injection volume, multiple injections were made at nearby depths or nearby sites within the target area in most cases. Total injection volumes, summed across all depths and all sites for each case, are listed in Table 1 (*M* = 3.1 µl, *SD* = 1.1 µl, n = 16). Following injections, the skull was sealed with dental cement before closing the scalp with surgical staples and reviving the rat.

**Table 1.** Injection sites and total injection volume at each site for all cases. mCherry and GFP refer to mCherry and GFP modified rabies viruses.

| Case | Virus | Inj site | Injection volume (uL) |
| --- | --- | --- | --- |
| R1630 | mCherry | AL | 1.2 |
| R1703 | mCherry | V1 | 3.6 |
|  | GFP | V1 | 3.6 |
| R1802 | mCherry | V1 | 2.4 |
|  | GFP | V1 | 2.4 |
| R1803 | GFP | LM | 1.2 |
| R1902 | mCherry | AM | 4.8 |
|  | GFP | PM | 4.8 |
| R1903 | mCherry | LP<sub>rm</sub> | 3.6 |
|  | GFP | LPI | 3.6 |
| R1905 | mCherry | LPI | 1.8 |
|  | GFP | LP<sub>rm</sub> | 1.8 |
| R1908 | mCherry | LP<sub>rm</sub> | 3.6 |
|  | GFP | LPI | 3.6 |
| R1909 | mCherry | LPI | 3.6 |
|  | GFP | LP<sub>rm</sub> | 3.6 |

Following a 7-14 day survival period, rats were deeply anesthetized with Euthasol and transcardially perfused first with saline, then with 4% paraformaldehyde in PBS. Brains were removed and cryoprotected in 30% sucrose for at least 24 hours, then sectioned coronally on a freezing microtome to 40 µm thickness, mounted on glass microscope slides, and coverslipped using polyvinyl alcohol mounting medium with 1,4-diazabicyclo-octane (PVA-DABCO, prepared in-house).

To assess corticothalamic connectivity, every fourth section was scanned using a fluorescent microscope (Axioplan 2, Zeiss, White Plains, NY) equipped with a 10x objective and motorized stage. Images were captured with a monochromatic low-noise CCD camera (Sensicam qe, PCO AG, Kelheim, Germany) and corrected for lamp misalignment by dividing each pixel by corresponding pixels in a flat field image acquired for each color channel. Corrected images were stitched using stage coordinates with regions of 10 overlapping pixels between images in which average pixel values were used. False colors were applied to each image before each brain section was counted manually for labeled cells. Neurons were identified based on the presence of the cell soma and dendrites. Fluorescently labeled neurons were then annotated by anatomical brain region based on the rat brain atlas by Paxinos and Watson (2013). Injection sites were identified in histology by tracks left by the glass micropipette (see Figure 3b). Image correction and stitching were performed in MATLAB (Mathworks, Natick, MA) using the multisection-imager toolbox (http://github.com/leoscholl/multisection-imager).

**Figure 3.**
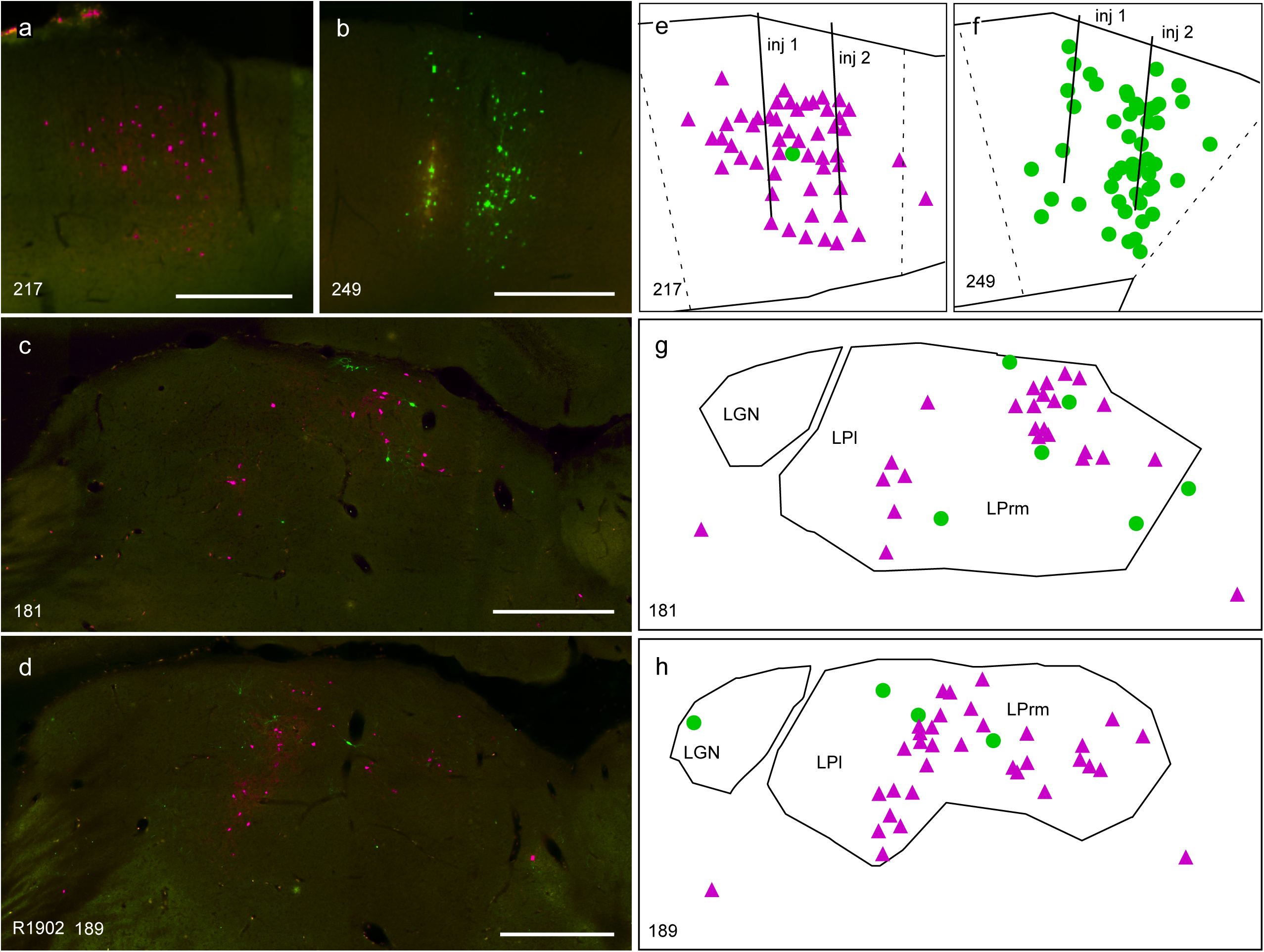
Representative example showing fluorescently labeled thalamic inputs to anterior and posterior medial V2 areas AM and PM after injections of mCherry (a and e; magenta) and GFP (b and f; green) modified rabies viruses in rat R1902. Thalamic labeling (c-h) reveals a large population of LPl and LPrm projections to V2. False-color stitched fluorescent images are shown in a-d. Reconstructions of the same sections are shown in e-h. Scale bars equal 500 µm.

Statistical significance was determined based on uncorrected cell counts. P values lower than 0.05 were considered significant for Student’s *t*-tests and two-way ANOVAs. All statistical analyses were carried out in MATLAB.

## Results

To assess the strength and size of input from LP to V1, we made four injections of the retrograde fluorescent-protein-expressing g-deleted rabies virus into V1 of two rats. In each case, substantial thalamic labeling was observed when calculated as a percentage of the total number of labeled neurons in each case (*M* = 11%, *SD* = 6%, n = 4).

In rat R1703, one large injection was made in anterior V1 (GFP; Figure 4a), and a second in posterior V1 (mCherry; Figure 4a). The resulting fluorescent labeling included a large number of cortical cells in and around the injection sites. In the thalamus, 30 labeled cells were found in LGN and only seven in LP across the two injections (see Table 2). Topographic organization of LGN labeling was observed between the two injection sites, with the posterior V1 injection labeling anterior LGN and vice versa, consistent with previous findings in the rat (Sauve & Gaillard, 2007). No such topography was seen for LP cells in this case, although it has been reported in mice (Roth et al., 2015; Juavinett et al., 2019).

**Figure 4.**
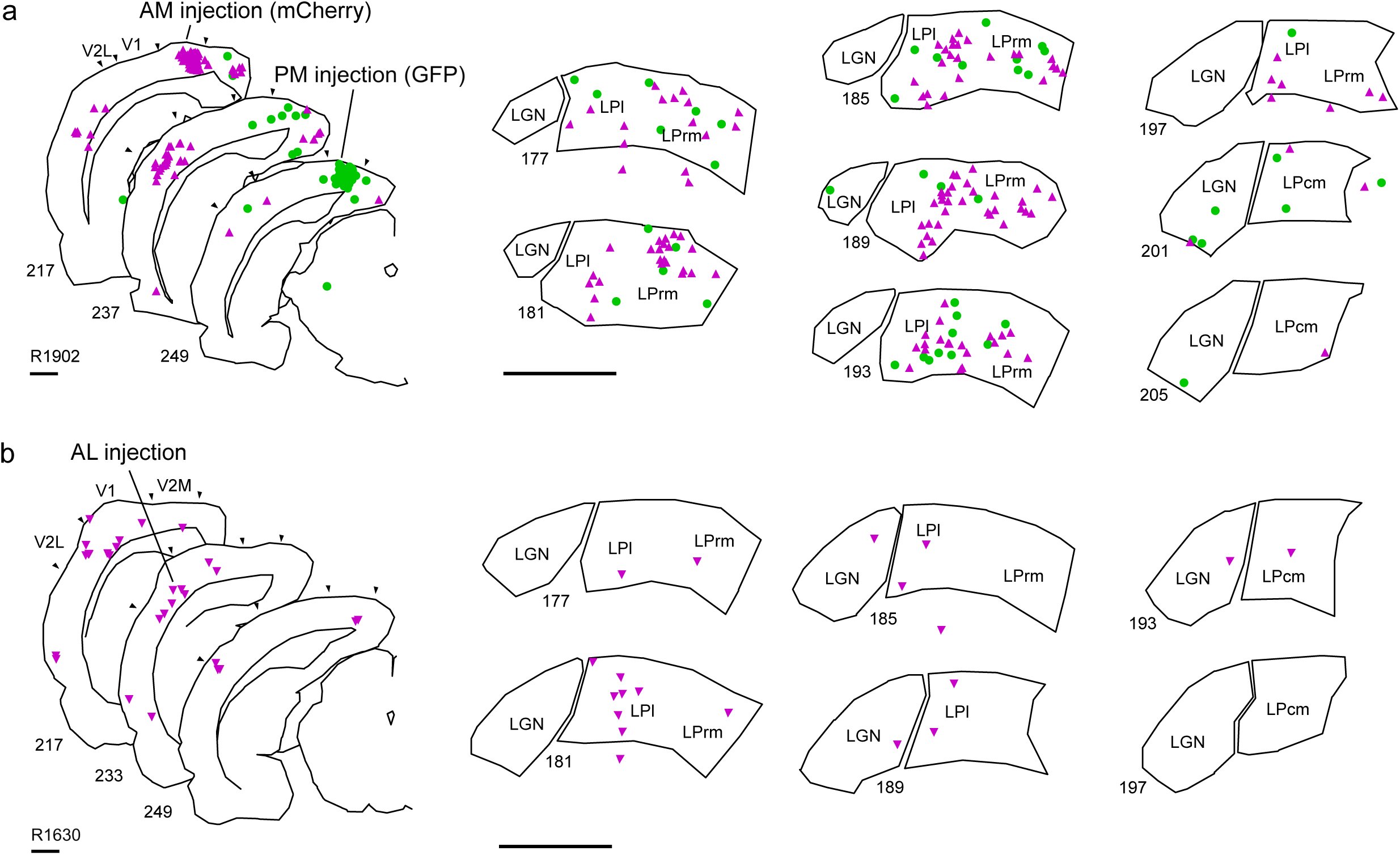
Reconstructions of fluorescently labeled neurons identified after retrograde RV injections in V1. In rat R1703 (a), the GFP virus was injected into the anterior portion of V1 and the mCherry virus was injected into posterior V1. In rat R1802 (b), three smaller injections were made along the anterior-posterior axis of V1 for each virus, covering a large portion of V1. In all cases, more labeled neurons were present in LGN than in pulvinar. Magenta triangles indicate mCherry fluorescence, green circles indicate GFP. Scale bars equal 1 mm.

**Table 2.** Percent of retrogradely labeled thalamic cells following cortical rabies virus injections.

| Rat | Virus | Target | No of cells | LP<sub>rm</sub> | LPI | LP<sub>cm</sub> | LGN |
| --- | --- | --- | --- | --- | --- | --- | --- |
| R1703 | GFP | V1 anterior | 16 | 6 | 13 | 0 | 81 |
|  | mCherry | V1 posterior | 21 | 0 | 19 | 0 | 81 |
| R1802 | mCherry | V1 | 34 | 6 | 9 | 6 | 79 |
|  | GFP | V1 | 26 | 4 | 8 | 4 | 85 |
|  |  |  | **Mean** | **4** | **12** | **2** | **82** |
| R1630 | mCherry | V2L (AL) | 19 | 11 | 63 | 11 | 16 |
| R1803 | GFP | V2L (LM) | 15 | 7 | 53 | 13 | 27 |
| R1902 | mCherry | V2M (AM) | 124 | 44 | 54 | 2 | 0 |
|  | GFP | V2M (PM) | 61 | 41 | 46 | 3 | 10 |
|  |  |  | **Mean** | **26** | **54** | **7** | **13** |
|  |  |  | **Total Mean** | **15** | **33** | **5** | **47** |

**Table 3.** Percent of retrogradely labeled cortical cells following LP injections

| Rat |  | Target | No of cells | V1 | V2 | medial V2 | lateral V2 |
| --- | --- | --- | --- | --- | --- | --- | --- |
| R1903 | mCherry | LP<sub>rm</sub> | 159 | 14 | 86 | 34 | 52 |
| R1905 | GFP | LP<sub>rm</sub> anterior | 35 | 17 | 83 | 77 | 6 |
| R1908 | mCherry | LP<sub>rm</sub> | 76 | 30 | 70 | 30 | 39 |
| R1909 | GFP | LP<sub>rm</sub> | 74 | 31 | 69 | 30 | 39 |
|  |  |  | **Mean** | **23** | **77** | **43** | **34** |
| R1903 | GFP | LPI | 425 | 32 | 68 | 21 | 47 |
| R1905 | mCherry | LPI anterior | 231 | 25 | 75 | 46 | 29 |
| R1908 | GFP | LPI | 109 | 28 | 72 | 18 | 54 |
| R1909 | mCherry | LPI | 131 | 35 | 65 | 37 | 28 |
|  |  |  | **Mean** | **30** | **70** | **31** | **40** |
|  |  |  | **Total Mean** | **26** | **74** | **37** | **37** |

In rat R1802, three small injections were made spanning anterior to posterior V1 for each virus (Figure 4b), in order to infect a large area of V1 axon terminals while keeping the total injection volume comparable to the previous injections. Labeled cells in LGN were spread along the anterior posterior axis in both cases. In total, 49 cells were labeled in LGN and 11 in LP across the six injections (see Table 2). Cells were found in all three LP subdivisions in both cases.

Across the four cases with V1 injections (see Table 2), there were significantly fewer inputs from LP to V1 compared to the number of inputs from LGN to V1 (*p* = 0.01, paired samples *t-*test). Infecting a large area of V1 on the possibility that LP input to V1 is sparse made no difference; there was one fifth as many cells in LP as in LGN in both R1703 and R1802.

In contrast to V1, injections in V2 labeled proportionately fewer neurons in LGN and many more neurons in LP. In rat R1902, the anterior and posterior medial V2 areas, AM and PM, were each injected with rabies virus (Figure 5a). The resulting fluorescence in the thalamus was primarily located within LP, 179 cells, compared to the LGN, 6 cells, indicating the injections were well contained outside V1. Between LP subdivisions, only four cells were labeled in LPcm, while 95 were labeled in LPl and 80 in LPrm (see Table 2 for summary). The cells in LPl and LPrm were not distributed throughout each subdivision, rather they were clustered along the medial margin of LPl and central LPrm (see Figure 5a).

**Figure 5.**
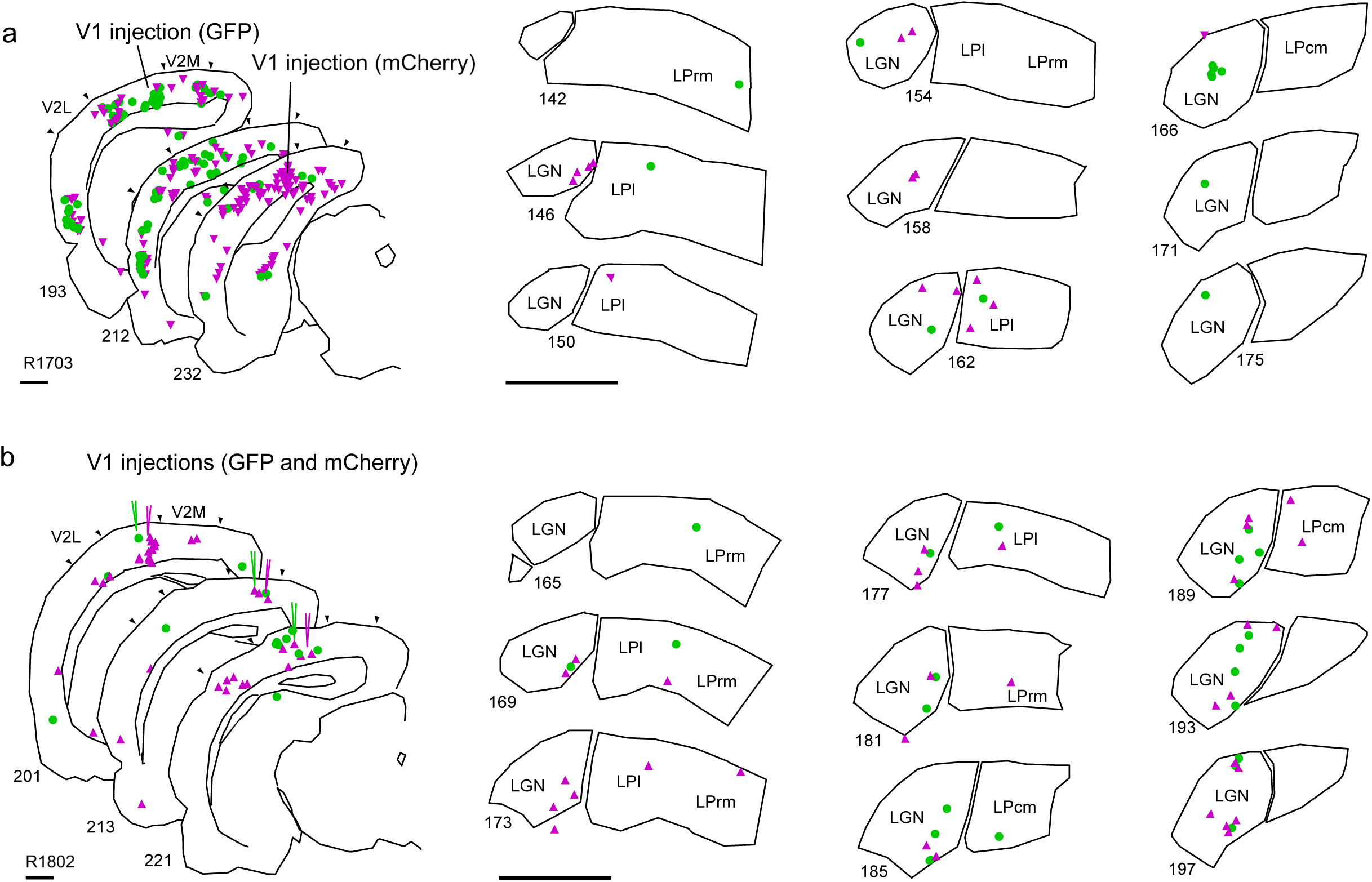
Reconstructed fluorescent labeling for rats with injections in V2 areas. Rat R1902 (a) was injected with mCherry virus targeting anteromedial V2 area (AM), and GFP virus targeting the posteromedial V2 area (PM). Labeling in the thalamus was strong in both LPrm and LPl, with only a few labeled cells in the lateral geniculate nucleus (LGN). Rat R1630 (b) was injected with mCherry virus targeting anterolateral V2 (AL), resulting in thalamic labeling mostly in LPl. Magenta triangles indicate mCherry fluorescence, green circles indicate GFP. Scale bars equal 1 mm.

Lateral V2 areas AL and LM were targeted with single injections into rats R1630 and R1803, respectively. These injections were three times smaller than the injections in R1902, yet thalamic labeling was also limited primarily to the LP in these cases, as 26 neurons were located in LP and 7 in the LGN. Between LP subdivisions, LPl had the majority of fluorescent cells in both cases, with 73% of LP cells being found within LPl in R1630 and 80% in R1803 (see Table 2).

Across all four cases with V2 injections, there was a significantly higher percentage of cells labeled in LP than in LGN (*p* < 0.01, paired samples *t-*test; see Table 2). Moreover, compared with V1 injections, V2 injections revealed that LP sends significantly more projections to V2 than it does to V1. The number of labeled cells in LP following V2 virus injections was significantly larger than the number following V1 injections (*F*(1,12) = 6.1, *p* = 0.03, *ANOVA*). Additionally, medial V2 areas AM and PM were both observed to receive more LP input than lateral V2 areas AL and LM, especially from LPrm. There was also some indication that lateral V2 areas receive less input from medial LP (n.s.), however there was no difference between overall input to V2 between the two rostral subdivisions of LP (*F*(1,12) = 0.28, *p* = 0.6, *ANOVA*).

Injections of modified rabies virus were also made into LPrm and LPl to retrogradely determine the strength of corticothalamic inputs from V1 and V2 to each pulvinar subdivision. Figure 6 illustrates three cases with injections targeting similar stereotaxic coordinates for LPrm and LPl. Rats R1903 (Figure 6a) and R1909 (Figure 6c) had large injections targeting central LPl and LPrm; rat R1909 (Figure 6b) had large injections targeting central LPl but more posterior LPrm; and rat R1905 (Figure 7) had small injections into anterior LPl and LPrm (see Table 1). Cortical labeling was assessed in each case to determine the relative strength of inputs from V1 and V2 to each rostral LP subdivision. In all cases, V2 labeling accounted for most of the cortical fluorescence. Rat R1903 additionally had significant reticular thalamic nucleus labeling following injection into LPl (Figure 6a), which is known to project to the pulvinar and many other neighboring thalamic nuclei (Lyon et al., 2010; Bourassa & Deschenes, 1995; Zikopoulos & Barbas, 2007; Hirsch et al., 2015). Also of note, rat R1909 had significant amygdala labeling following injection into LPrm, but this is perhaps due to leakage of virus into the hippocampus directly dorsal to LP.

**Figure 6.**
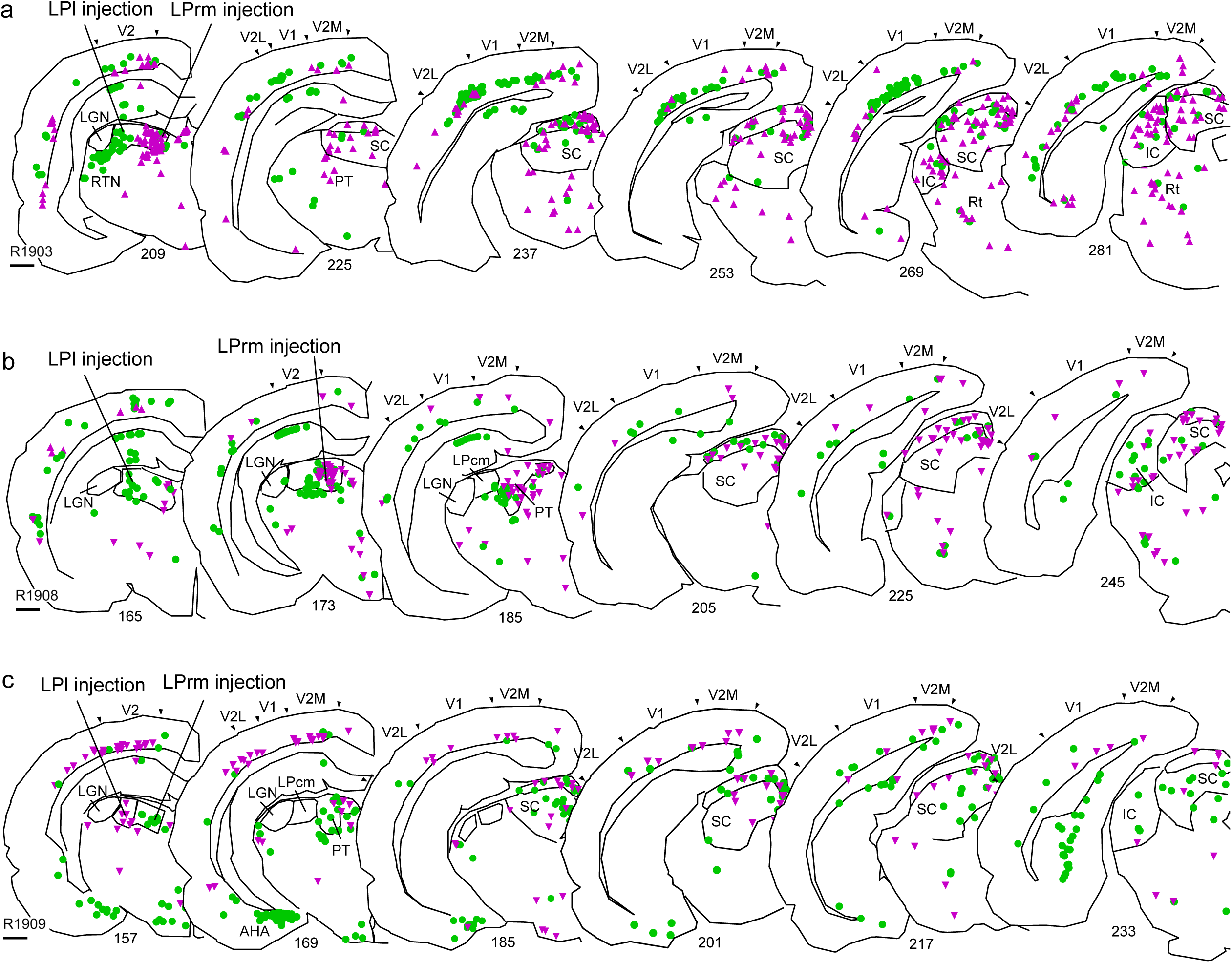
Reconstructions of fluorescently labeled neurons identified following rabies injections in LPrm and LPl in three rats. Each LPl injection and each LPrm injection was targeted to the same stereotaxic coordinates and used the same volume, with the exception of rat R1908 (b), where more posterior LPrm was targeted to avoid a blood vessel. The superficial layers of the superior colliculus are demarcated. Magenta triangles indicate mCherry fluorescence, green circles indicate GFP. IC inferior colliculus, RTN reticular thalamic nucleus, PT pretectal nucleus, Rt reticular formation. AHA amygdalohippocampal area. Scale bars equal 1 mm.

**Figure 7.**
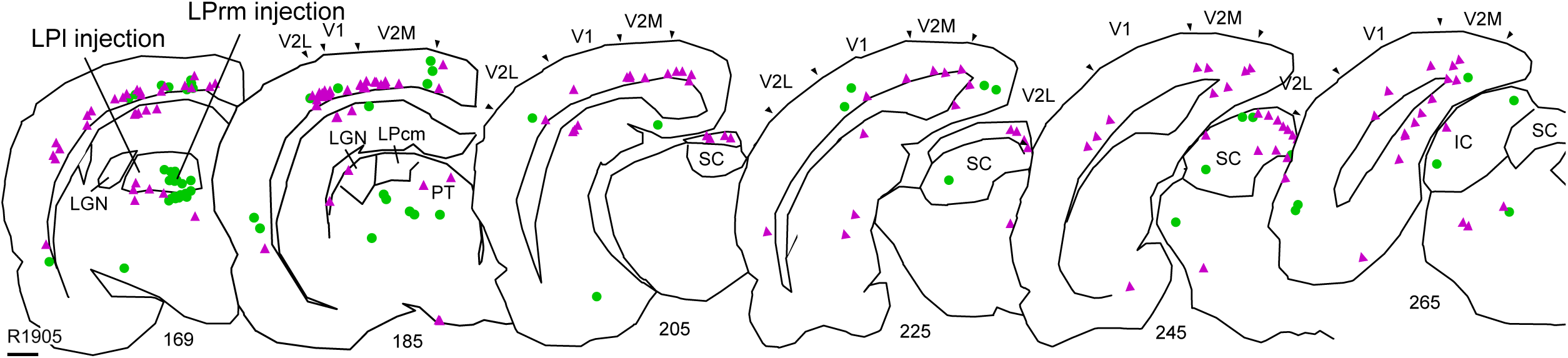
Reconstruction of rat R1905, in which retrograde injections were made in the rostral portions of LPl and LPrm. Very little labeling is apparent in pretectum (PT) and superior colliculus (SC) following GFP virus injection into LPrm in this case (green circles). Injection of mCherry virus (magenta triangles) yielded similar results to previous injections. Scale bar equals 1 mm.

Across all eight LP injections, there were a similar number of neurons labeled in SC as in visual cortex (*p* = 0.3, paired samples *t-*test). Since rostral LP receives most of its input from cortex, and caudal LP receives most of its input from SC (in rats: Mason and Groos, 1980, Takahashi, 1985, Masterson, 2009; in mice: Bennett, et al., 2019, Juavinett et al., 2019), this is a good indication that caudal and rostral LP in this study were infected at similar rates.

A significantly higher number of cells were labeled in V2 areas than in V1 (*F*(1,12) = 5.39, *p* = 0.04, *ANOVA*). Furthermore, injections in lateral LP led to significantly more labeling in visual cortex than injections in medial LP (*F*(1,12) = 5.35, *p* = 0.04, *ANOVA*), implying that rat pulvinar receives differential visual cortical inputs along its medial-lateral axis.

## Discussion

The purpose of this study was to compare the connections of V1 and higher visual cortex with the pulvinar as well as to determine any anatomical differences between lateral and medial pulvinar subdivisions in the rat. We demonstrated that the projections between pulvinar and V2 in rats are more frequent than those between pulvinar and V1, suggesting that the pulvinar has a greater influence on activity in higher visual cortex than in V1. We also observed differences in the number of inputs and outputs of LP subdivisions, with the lateral portion of the pulvinar having a stronger connection to visual cortex, for both V2 and V1. Together, these results provide a basis for understanding to which visual networks the rat pulvinar contributes.

In previous studies of the pulvinar, projections have been identified to both V1 and higher visual cortices, but no attempt to quantify these cells has been made. However, evidence consistent with our findings does exist. In rats, anterograde injections in LP labeled both V1 and V2, but V1 labeling was much more sparse, with many more fibers being labeled in both medial and lateral V2 (Nakamura et al., 2015, see their Figure 7). In mouse, retrograde injections in V1 produced fluorescent labeling that was confined to small regions within LP, whereas injections in other visual cortical areas led to larger and brighter patches of fluorescent labeling (Juavinett et al., 2019, see their Figure 4). Although qualitative, these previous findings are consistent with the difference in projection strength we observed in the present study.

In other species, there is also a trend in the existing literature of denser and more numerous projections from pulvinar to higher visual cortex compared to projections from pulvinar to V1. In squirrels, a highly visual rodent, dense pulvinar labeling was observed following retrograde tracer injections into the temporal posterior area and visual area 19, but very few cells were labeled following V1 (area 17) injection (Robson & Hall, 1977, see their Figures 15-17). Tree shrew, a close primate relative, exhibits a similar pattern of connectivity following retrograde tracer injections in V1 and V2; only sparse, topographic labeling was observed in caudal pulvinar following V1 injection, whereas V2 injection was followed by denser labeling spread across large regions of caudal and ventral pulvinar (Lyon et al., 2003, see their Figure 2). Similarly, in monkeys, retrograde injections in V1 of macaque labeled very few cells in pulvinar compared to injections in V2 in marmoset (Kaas & Lyon, 2007, see their Figures 6-7). Although this pattern seems to have gone largely unnoticed, it is present in all of the anatomical literature we have come across.

Given that several high profile studies have found significant functional effects of pulvinar on V1 in other species (Purushothaman, et al., 2012, Roth et al., 2015, Sun et al., 2016), it is tempting to assume that there must be a significant population of pulvinar neurons projecting directly to V1 in these animal models. However, our results indicate that higher visual cortex receives significantly more input from the pulvinar than does V1, thus pulvinar would likely have a bigger impact on visual responses there, and the gating of V1 responses by pulvinar could be explained by indirect modulation via higher visual cortex. This view is consistent with the results of Tohmi et al. (2014), who showed higher visual areas in mice behave more like V1 when the extrageniculate pathway is damaged by superior colliculus lesions. In addition, Zhou et al. (2016) found that deactivation of the ventrolateral pulvinar in monkeys led to inactivity in V4, without evidence for change in V1 activity, implying a direct influence of pulvinar on V4. Nevertheless, as we and others observe some pulvinar projections to V1, direct effects on V1 are not out of the question.

In primates, the pulvinar is divided by its connectivity with cortical areas into a dorsal-ventral stream classification for visually guided actions and object vision, respectively. The posterior and caudomedial inferior pulvinar (PIp and PIcm) as well as the medial inferior pulvinar (PIm) are associated with the dorsal stream because they project to dorsal stream areas such as the middle temporal visual area (MT), whereas the caudolateral inferior (PIcl) portion and lateral (PL) portions of the pulvinar are associated with the ventral stream because of their projections to early visual areas and inferior temporal cortex (Kaas and Lyon, 2007). As illustrated in Figure 8, SC and visual cortex input to pulvinar is highly conserved across species, with regions of dense bilateral input from SC, ipsilateral input from SC, and cortical input only. These include rat (Takahashi, 1985; Mason & Groos, 1980), mouse (Zhou et al., 2017), gray squirrel (Baldwin et al., 2011), tree shrew (Lyon et al., 2003), and primate (Baldwin et al., 2013). Given that they share anatomical features with primates, rodents might be a suitable model animal for studying pulvinar contributions to the two visual streams, as there is growing evidence for such a classification in rodents (see Figure 9; Wang et al., 2011; Glickfeld, 2014; Nishio et al., 2018). However, previous studies have shown that tectorecipient LPcm in rats sends most of its projections to ventral stream areas such as postrhinal cortex (POR) and temporal association cortex (Nakamura et al., 2015), and our results show that LPrm and LPl are well connected with dorsal stream areas AM and PM, opposite to the pattern of tecto- and striate-recipient zones of the pulvinar in monkeys. In mice, however, LPrm was observed to receive significant input from POR, and LPl significant input from LM (Juavinett et al., 2019), so the distinctions may not be so clear cut.

**Figure 8.**
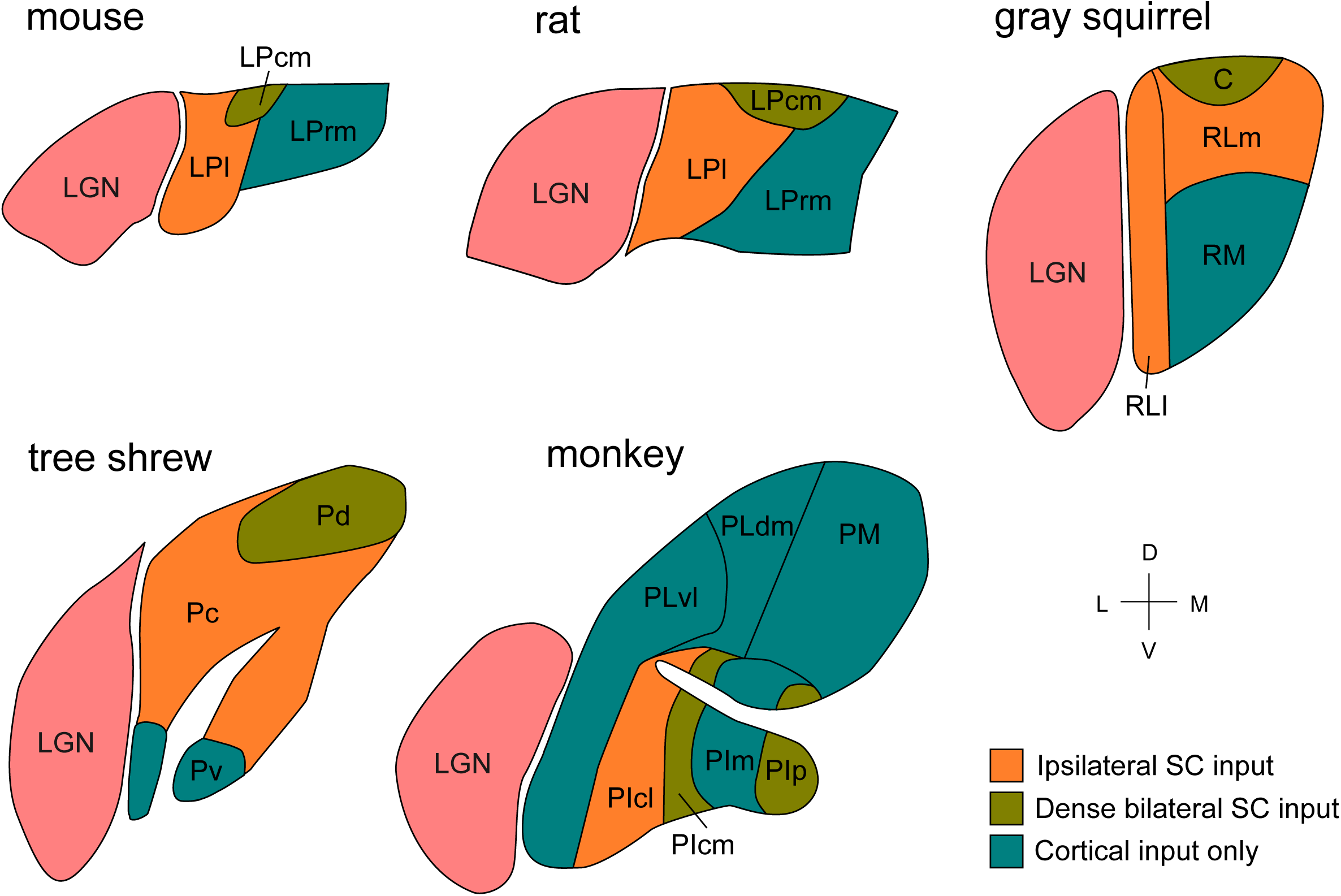
Conserved pulvinar input scheme from SC and visual cortex across species. Schematic diagrams of the pulvinar in mouse, rat, gray squirrel, tree shrew, and monkey are shown. Areas with dense bilateral input from SC are shown in green, areas with ipsilateral SC input are shown in orange, and cortical recipient areas are shown in blue. Adapted from Lyon et al. (2003) and Roth et al. (2017).

**Figure 9.**
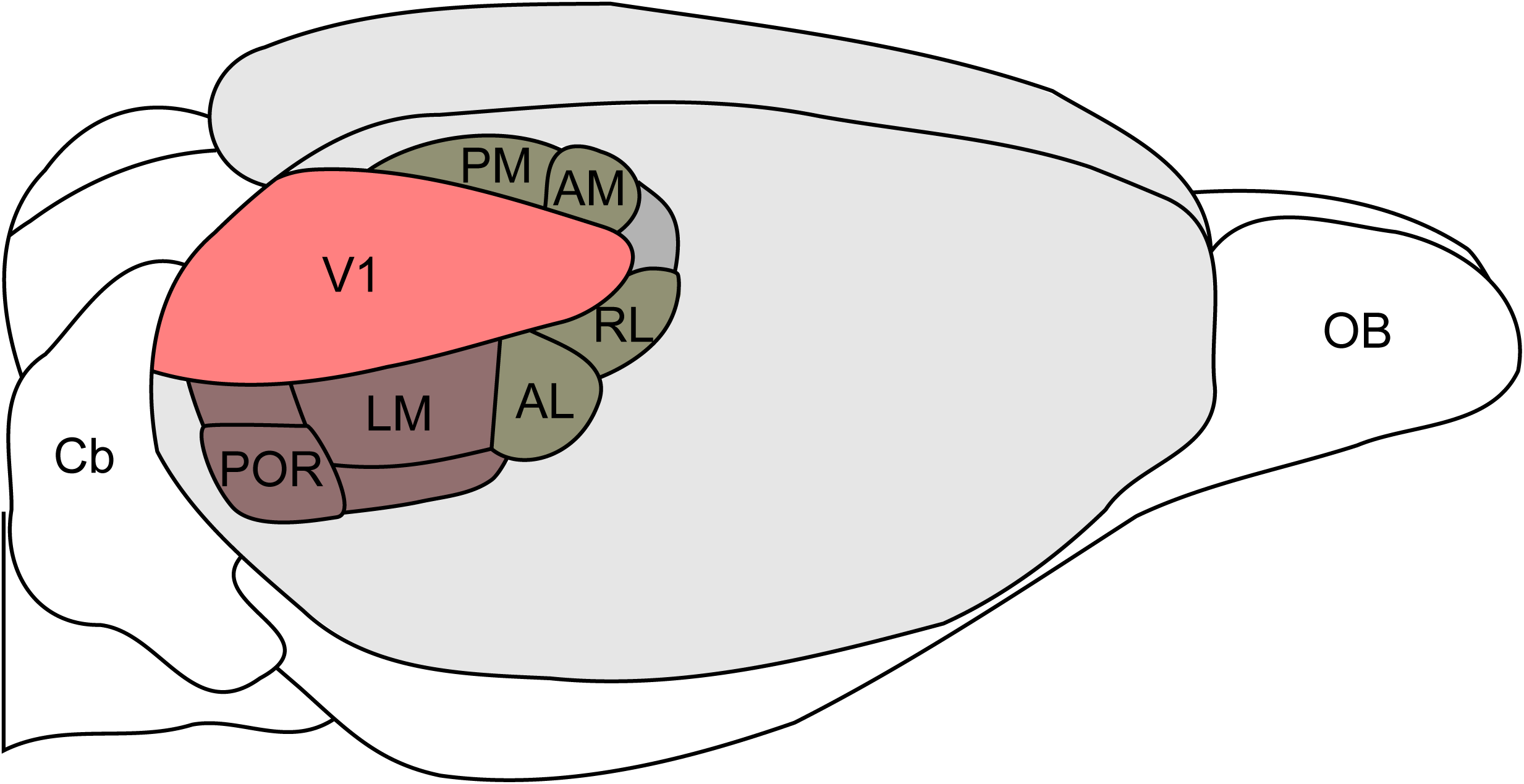
Summary of dorsal and ventral stream areas in the visual cortex of the rat. Modified from Sereno et al. (1991) based on Espinoza et al (1983), Glickfeld et al. (2014), and Wang et al. (2007). Cb cerebellum, OB olfactory bulb.

In monkeys, disruption of the dorsolateral pulvinar has been shown to cause deficits to spatial attention (Wilke et al., 2010). It is unclear which pathways the pulvinar influences, and whether it generates salience-based signals or receives top-down influence from other cortical areas. If pulvinar is responsible for modulating cortex based on salience cues, it seems unlikely that these salience maps would follow the hierarchy of visual areas as suggested by Zhang et al. (2012), since pulvinar is only weakly connected to V1. Instead, V1 salience maps might derive from LGN, while pulvinar modulates higher-level features such as motion or texture in other cortical areas (Schiller, 1993; Saalmann et al., 2012; Perry & Fallah, 2014). If, on the other hand, pulvinar receives attention signals from higher visual cortex, such as by V4 as demonstrated by Zhou et al. (2016), then reciprocal connections with these areas, but not V1, would be sufficient. Further research will be needed to evaluate pulvinar’s functional role in attention.

## Acknowledgements

This work was supported by the Whitehall Foundation #2014-08-100 (DCL) and the National Eye Institute R01EY024890 (DCL).

